# Renal arterial dysfunction, impaired pressure natriuresis and salt-sensitivity in a mouse model of Cushing syndrome

**DOI:** 10.1101/2024.12.11.625204

**Authors:** Hannah M Costello, Céline Grenier, Jessica R Ivy, Natalie K. Jones, Kevin Stewart, Megan C Holmes, Dawn E.W. Livingstone, Neeraj Dhaun, Matthew A Bailey

**Affiliations:** Edinburgh Kidney, British Heart Foundation Centre for Cardiovascular Science, The University of Edinburgh, United Kingdom, EH16 4TJ

**Author notes:** For Correspondence: Matthew Bailey, PhD, FRSB.

## Abstract

**Background:** Cushing Syndrome arises from endogenous overproduction of ACTH by a tumour or is acquired through chronic exposure to glucocorticoid medication. Hypertension is a major complication, increasing cardiovascular risk, but underlying mechanisms are not clearly defined.

**Methods:** We infused male C57BL/6J mice with ACTH or vehicle for 14-21 days, inducing cardinal features of Cushing Syndrome. The renal pressure natriuresis response was measured under anesthesia and ex vivo artery function assessed by myography.

**Results:** ACTH infusion blunted the natriuretic and diuretic responses to incremental increases in blood pressure. Renal hemodynamics did not change with blood pressure in controls, but renal autoregulation was impaired in ACTH mice. The *ex vivo* contractile response of the renal artery to phenylephrine was diminished in Cushing Syndrome mice, as was endothelium-dependent and endothelium-independent relaxation. On 0.3% sodium diet, there was no evidence of sodium retention in ACTH-treated mice but the diurnal rhythm of sodium excretion was abnormal and mice had non-dipping BP. The Cushing Syndrome model also displayed enhanced salt preference and amplified salt-sensitive blood pressure.

**Conclusion:** Cushing Syndrome induces a cluster of phenotypes impacting sodium homeostasis and blood pressure regulation. Hypertension, salt-sensitivity and non-dipping blood pressure are important cardiovascular risk factors, and, beyond Cushing Syndrome, our findings are relevant to obesity and the metabolic syndrome, in which tissue glucocorticoid homeostasis is abnormal.

## INTRODUCTION

Cushing disease, caused by an adrenocorticotropic hormone (ACTH)-secreting pituitary tumour, is the most common endogenous form of Cushing Syndrome. It causes glucocorticoid excess and substantially increases cardiovascular risk^1^. The condition is rare but Cushing Syndrome arising from prescribed glucocorticoid therapy has a broader public health impact. Synthetic glucocorticoids are front line treatments for many inflammatory conditions; ∼2% of the adult population receive glucocorticoid prescriptions and there is an upward trend in prescriptions for sustained therapy in conditions such as chronic obstructive pulmonary disease^2^. About 60% of people receiving prolonged glucocorticoid therapy develop iatrogenic Cushing syndrome and acquire the burden of increased cardiovascular risk^3^.

The causes of cardiovascular disease are not fully defined but high blood pressure (BP) and loss of the physiological nocturnal BP dip are common complications of Cushing Syndrome. Hypertensive mechanisms are complex due to pleiotropic effects of glucocorticoids on BP regulatory systems, including arterial vascular tone and renal sodium excretion^4,5^. Abnormal activation of the renin-angiotensin-aldosterone system (RAAS) is well documented and the pressor response to angiotensin II is amplified^6^. Glucocorticoid excess may also induce sympathetic hyperactivity and an exaggerated pressor response to noradrenalin has been reported in patients with Cushing Disease^7^. Indeed, sustained glucocorticoid excess impacts many vasoregulatory systems and vasoconstriction is augmented^4^.

Increased peripheral resistance is a significant factor for the onset and maintenance of arterial hypertension in Cushing Syndrome. Whether abnormal sodium handling by the kidney contributes is less certain. Increased renal sympathetic nerve activity and elevated angiotensin II promote sodium retention, as does hyperactivation of the mineralocorticoid receptor (MR) by glucocorticoids when 11β hydroxysteroid dehydrogenase type 2 (11βHSD2) is saturated^8^. Additionally, activation of the glucocorticoid receptor (GR) stimulates sodium reabsorbing mechanisms in the proximal tubule^9^, thick limb of Henle^10^, and distal convoluted tubule^11^, and GR activation may also modulate epithelial sodium channel (ENaC) activity in the collecting duct^12,13^, although the physiological role is less certain^14^. The increased abundance of phosphorylated NKCC2 and NCC in urinary vesicles from patients with Cushing Syndrome is consistent with induction of epithelial sodium transport^15^. In healthy humans, a 5-day infusion of either ACTH or cortisol reduces sodium excretion and expands plasma volume^16^. In a mouse model of Cushing Syndrome, ENaC activity is increased through MR and GR pathways^17^. However, other transporters are down-regulated and overall ACTH-infused mice are in negative sodium balance^18^. Nevertheless, dietary sodium depletion prevented hypertension in Cushing Syndrome mice^17^, and such salt-sensitivity is indicative of a renal phenotype^19^.

The kidney determines long-term stability of BP by controlling extracellular fluid volume (ECFV) through the pressure-natriuresis response^20,21^. This response has both renal vascular/hemodynamic and tubular/epithelial components that can be influenced by glucocorticoid excess and we hypothesized impairment of these in Cushing Syndrome. We tested this hypothesis in male C57BL/6JCrl mice infused with ACTH for 14-21 days, which we have previously shown captures the endocrine, metabolic, and cardiovascular features of Cushing Syndrome^22,23^.

## METHODS

Experiments described were performed between September 2018 and September 2021 in the Centre for Cardiovascular Science at the University of Edinburgh. Adult male C57BL/6 mice (age 10-12 weeks) were commercially sourced (Charles River, UK) and maintained in our animal unit under controlled conditions (temperature (24 ± 1⁰C), humidity (50 ± 10% humidity) and light/dark cycle (lights on 7am to 7pm local time)). Mice had ad libitum access to water and commercial rodent chow. The control diet contained 0.3% sodium and 0.7% potassium by weight [RM1, SDS Diets, United Kingdom (UK)]; the high salt diet contained 3% sodium and 0.6% potassium by weight (RM 3% Na+ SY, SDS Diets, UK). Mice were randomized into treatment groups and experiments performed with a single blinding to group allocations. All procedures conformed to the guidelines from Directive 2010/63/EU of the European Parliament on the protection of animals used for scientific purposes. Experiments were performed in accordance with the UK Animals (Scientific Procedures) Act under a UK Home Office Project Licence to the lead investigator (MAB) following ethical review by the University.

### ACTH-treatment

Chronic administration was achieved by subcutaneous osmotic minipumps (Alzet, models 2002/2004, Charles River UK) which infused ACTH (2.5 μg/day, adrenocorticotropic hormone fragment 1-24 human, rat; Sigma, MO, USA) for periods of up to 28 days as stated for each experiment. Pumps were primed in sterile saline at 37°C per manufacturer’s instructions and implanted aseptically under isoflurane anesthesia (4% induction; 2% maintenance), with postoperative buprenorphine administration (0.1 mg/kg; Vetergesic; sc) for analgesia and 0.9% sterile saline (0.5mL; IP).

### Characterization of Cushing Syndrome features

Body weight and gonadal fat pad weight normalized to body weight were measured following 14 days of ACTH or vehicle infusion. Dissected left adrenal glands from control and ACTH mice were fixed in 4% paraformaldehyde for 24 hours at room temperature before being transferred to 70% ethanol. Samples were dehydrated, using an ethanol series, followed by xylene and embedded into paraffin blocks. Fixed adrenal glands were sectioned (5 µm), stained with hematoxylin and eosin (H&E) and examined by light microscopy and photographed by an AxioScan slide scanner (Zeiss, Cambridge, UK). Adrenal gland cross-sectional area was calculated (QuPath). The adrenal glands were analysed in a randomized order by an observer blinded to the treatment groups.

Corticosterone was measured in plasma samples taken by tail venesection from the same mouse at the diurnal nadir (7am) and peak (7pm) in mice and measured by commercial ELISA kit (ADI-900-907; Enzo Life Sciences, UK). The cross-reactivity for this kit is 100% for corticosterone, 21.3% 11-deoxycorticosterone, 21.0% desoxycorticosterone, 0.46% progesterone, 0.31% testosterone, 0.28% tetrahydrocorticosterone, 0.18% aldosterone, 0.046% cortisol, and <0.03% for pregnenolone, estradiol, cortisone, 11-dehydrocorticosterone acetate. For aldosterone, trunk blood was collected onto ice following decapitation at either 7-8am or 7-8pm. Plasma aldosterone was measured by commercial ELISA (ADI-900-173; Enzo Life Sciences). The cross-reactivity for this kit is 100% for aldosterone, 0.3% 11-deoxycorticosterone, 0.19% corticosterone, 0.2% progesterone, and <0.001% for cortisol, dihydrotestosterone, estradiol, and testosterone.

### Evaluation of renal function

#### Acute pressure natriuresis response

Pressure natriuresis experiments were carried out in control mice and mice infused with ACTH for 21 days. Under intraperitoneal barbiturate non-recovery anesthesia (Inactin, 120 mg/kg; Sigma), the right jugular vein was cannulated for intravenous infusion of physiological saline (100 mM NaCl, 15 mM NaHCO3, 5 mM KCl, 0.25% fluorescein isothiocyanate (FITC)-inulin (Sigma), 2% bovine serum albumin (Sigma), pH 7.4) to allow for calculation of glomerular filtration rate (GFR) and for maintenance of anesthesia. A tracheotomy was performed to relieve obstruction to breathing. The right carotid artery was cannulated for real-time mean arterial BP measurements (PowerLab, ADInstruments, Oxford, UK) and for intermittent blood collections. The bladder was catheterized to collect urine to measure urine flow rate and electrolyte excretion. Following a 30-minute equilibrium period, blood samples were collected for plasma (P1-P4) before and after 3 urine collections (U1-U3). U1 represented a 30-minute baseline period. The 1^st^ pressure ramp was then induced by sequential arterial ligation of the mesenteric and coeliac arteries (U2, 20 minute), followed by a 2^nd^ pressure ramp via distal aorta ligation (U3, 20 minute). BP was measured throughout baseline, 1^st^ and 2^nd^ pressure ramp, as well as urine flow rate, sodium excretion and GFR. Urinary sodium concentrations were analysed using SpotChem EL SE-1520 analyser (Arkray, Kyoto, Japan). GFR was calculated as the ratio of urinary to plasma FITC-inulin concentrations multiplied by the urine flow rate. The pressure natriuresis and diuresis response and change in fractional sodium excretion were also calculated.

#### Renal blood flow

Renal blood flow was measured on the left renal artery in a sub-group of control and ACTH-treated mice who went through the pressure natriuresis protocol using a pulse-wave Doppler ultrasound probe (Transonic, NY, USA). Renal vascular resistance was calculated dividing mean arterial BP by renal blood flow.

#### Metabolic cages

In a separate cohort of mice, sodium and fluid balance were assessed in conscious mice single-housed in metabolic cages for 24 hours at baseline and 14 days post-ACTH infusion. Daily food intake and water intake were measured. Urine was collected over consecutive 12-hour periods. Electrolytes were measured in urine and plasma using the SpotChem analyser (Arkay). Osmolality was measured by freezing-point depression (Camlab, Cambridge, UK). Plasma copeptin concentration measured by commercial (Cloud-Clone Corp., TX, USA).

### Ex vivo vascular function

Following decapitation between 7 am and 8 am, dissected renal and second order mesenteric arteries from control and ACTH-treated (14 days) mice were cleaned of adherent perivascular fat in physiological salt solution (PSS; 119.0 mM NaCl, 3.7 mM KCl, 2.5 mM CaCl2, 1.2 mM MgSO4, 25.0 mM NaHCO3, 1.2 mM KH2PO4, 27.0 μM EDTA, and 5.5 mM D-glucose) on ice and mounted on a wire myograph (DMT, Denmark). Each vessel mounted on the myograph was normalized using the DMT Normalization Module on LabChart (ADInstruments) to calculate and set the optimal pre-tension conditions by determining its internal circumference (µm). Vessels were equilibrated for 30 minutes in PSS and perfused with 95% O_2_ and 5% CO_2_ at 37°C. High potassium PSS (KPSS; 125 mM KCl) was used as a depolarizing contraction and maximal contraction was determined at the start and end of the experiment. Vascular reactivity was assessed by generating cumulative concentration-response curves to phenylephrine (1 nM to 30 μM PE; Sigma), acetylcholine (1nM to 30 μM ACh; Sigma) and sodium nitroprusside (1nM to 30 μM SNP; Sigma), separated by 30-minute washout periods. PE responses were calculated as a percentage of maximal KPSS response. ACh and SNP responses were calculated as a percentage of PE-induced pre-constriction.

### Quantitative PCR

Renal and second order mesenteric arteries were dissected and cleaned of perivascular fat from control and ACTH-treated (14 days) mice following decapitation. RNA was isolated from pooled vessels (4 renal arteries or 2 mesenteric arteries) using the RNeasy Micro kit (Qiagen, USA), quantified by the NanoDrop-1000 (Thermo Fisher Scientific, MA, USA) and RNA integrity assessed using automated electrophoresis (Agilent, CA, USA). cDNA (500ng) was generated using high-capacity RNA-to-cDNA kit (Thermo Fisher Scientific). Renal mesenteric artery abundance of *Adra1a* (encoding α1A adrenoreceptor), *Nos3* (endothelial nitric oxide synthase), *Prkg1* (encoding α- and β-isoforms of soluble cyclic GMP-dependent protein kinase 1), *Fkbp5* (FK506 binding protein 5), *Nr3c1 (*glucocorticoid receptor), *Hsd11b1 (*11β hydroxysteroid dehydrogenase type 1), *Nr3c2 (*mineralocorticoid receptor), *Hsd11b2 (*11β hydroxysteroid dehydrogenase type 2)*, Eln* (elastin), *Emilin-1, Vegfr2* and *Tgf-b1* were measured by quantitative RT-PCR using the Universal Probe Library (Roche, Switzerland). Primer sequences and probes numbers are given in **Supplemental table 1**. Triplicates of each sample and standard curve were run on the LightCycler 480 (Roche). A panel of appropriate reference genes, *Hprt*, *Rn18s*, *Actb* and *Tbp*, were selected for renal and mesenteric arteries on the basis that they did not differ between treatment groups. Data were then log transformed to ensure symmetry of the data set and normality tests were conducted. RT-qPCR data were handled in accordance with the MIQE guidelines.

### BP measurement by radiotelemetry

Under isoflurane anesthesia (4% induction; 2% maintenance), a PA-C10 radiotelemetry device (Data Sciences International, MN, USA) was implanted into a subcutaneous pocket with the sensor secured in the left carotid artery. Ocular lubricant (Lactri-lube) was applied to prevent eye dry out during surgery and buprenorphone (0.1 mg/kg; Vetergesic; sc) and sterile saline (0.5mL of 0.9% sterile saline IP) were administered postoperatively and mice were placed in a heated cage until mobility was fully regained. For 48 hours post-surgery, mice had *ad libitum* access to Vetergesic jelly and mash diet (RM1 diet soaked in drinking water). BP and locomotor activity recordings started 7 days after device implant, with data acquisition at 1Hz for a 5-minute period every 30 minutes. In one cohort of mice, baseline was recorded for 3 days before 21 days of ACTH infusion. Twenty-four-hour averages of systolic and diastolic BP were calculated, as well as BP dip (day BP-night BP) and locomotor dip. In a separate cohort of mice, the BP response to matched high salt intake salt-sensitivity before and after ACTH-infusion was assessed. Systolic and diastolic BP were measured following 7 days control diet, 7 days high salt diet, 5 days washout period, 12 days ACTH infusion on control diet before transition to a high salt diet for 16 days. Average steady state BP, average diurnal variation and BP dip of systolic and diastolic BP were assessed.

### Salt preference

In a cohort of control and ACTH-treated mice, salt preference was assessed using a two-bottle preference test pre- and post-minipump implantation, where the first drinking bottle was deionised water and the second was saline solution (1%). Bottles were alternated every day to avoid side preference.

### Statistical analysis

Data are presented as mean ± standard deviation (SD). Prior to analysis, the distribution of data was assessed using the Shapiro-Wilk normality test. Data were defined as normally distributed when *p* >0.05. Therefore, parametric analysis, including Student’s *t* test or ANOVA with Holm-Šidák’s *post hoc* test, was carried out, where appropriate, using GraphPad Prism 10.0 Software. A value of *P*<0.05 was considered significant. The number of biological replicates (*n*) and statistical analysis used are stated in the figure legends.

## RESULTS

Sustained infusion of ACTH reproduced cardinal features of human Cushing Syndrome with increases in gonadal fat deposits and adrenal gland size, alongside increases in circulating corticosterone and aldosterone (**Supplemental figure 1**). Additionally, the normal diurnal variation in circulating corticosterone was lost (ANOVA main effects: Group *P*<0.0001, diurnal *P*=0.121; interaction *P*=0.002), that of aldosterone maintained (ANOVA main effects: Group: *P*=0.024; diurnal *P*<0.0001; interaction *P*=0.801).

### ACTH infusion impairs the acute pressure natriuresis response

After 21 days of ACTH or vehicle infusion, mice were anesthetised for measurement of BP and renal function (**Figure 1A**). Baseline BP was ∼15 mmHg higher in ACTH-infused mice (**Figure 1B**) but sodium excretion (**Figure 1C**) and GFR (ACTH: 0.35±0.06 ml/min *vs* Control: 0.34±0.04 ml/min) were similar to vehicle-treated animals. To assess the pressure-natriuresis response, BP was acutely raised through sequential arterial ligation. This increased sodium excretion and urine flow rate in all mice. However, the natriuretic and diuretic response to increased pressure was significantly attenuated in the ACTH group (**Figures 1D & 1E**). Notably, at peak BP, GFR was not increased (Control: 0.32±0.11 ml/min *vs* ACTH: 0.38±0.07 ml/min) in either group of mice. Sodium excretion was plotted against BP to generate the pressure-natriuresis curve, which was significantly different in ACTH-treated mice (**Supplemental figure 2**). At peak BP, fractional sodium excretion was lower in ACTH-treated mice (**Supplemental figure 2C**), indicating impaired downregulation of tubule sodium reabsorption with rising BP.

**Figure 1.**
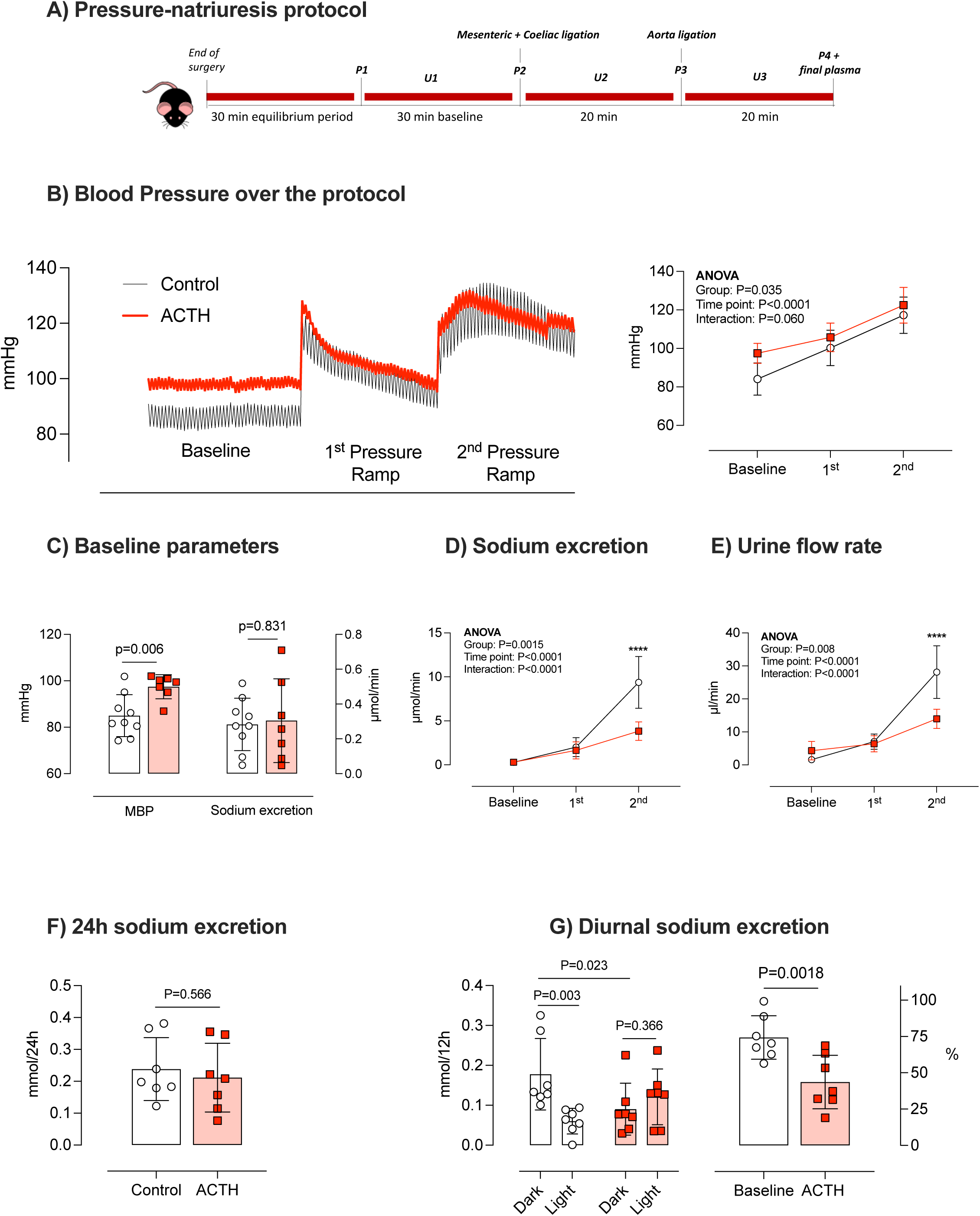
Renal excretion. Mice were infused with ACTH (red square; n=7) or saline vehicle (controls; open circle; n=9) for 21 days and were then anesthetised to measure the pressure natriuresis response. **A)** mice were surgically prepared to allow urine collection at baseline (U1) and over two consecutive periods (U2, U3) in which BP was acutely increased by arterial ligation. Plasma samples (P) were taken to allow measurement of FITC-inulin, used to calculate GFR and a final 0.5 ml sample to measure electrolytes. **B)** An example of an arterial blood pressure profile through the protocol. **C)** baseline mean arterial blood pressure (MBP) and sodium excretion, showing individual data points and group mean±SD. Statistical comparisons were made by unpaired t-test, with two-tailed P values shown. Group mean±SD for **D)** sodium excretion and **E)** urine flow rate in the three experimental periods. Statistical comparisons were made using two-way ANOVA with Holm-Šidák used for planned comparisons: ***=*P*<0.0001 between Control and ACTH groups at a given time point. In a separate experiment urine collections were made in n=7 mice before and after ACTH infusion. Two consecutive 12-hour collections were made to encompass the period of activity (Dark phase: subjective night) and inactivity (Light phase, subjective day). Collections were used to measure **F)** 24-hour sodium excretion and G) sodium excretion in dark and light phases, expressed in the right panel as a percentage of the total sodium output excreted in the dark phase. Statistical comparisons were made by two-way ANOVA with Holm-Šidák used for planned comparisons and by unpaired t-test.

To understand the impact on sodium and fluid homeostasis, a different group of mice was individually housed in metabolic cages for 24 hours at baseline and again after ACTH infusion. ACTH excess induced hypokalemia. These mice also had hypernatremia and increased plasma osmolarity and increased plasma copeptin, which are all indicative of volume contraction (**Supplemental table 1**). Total sodium excretion over 24 hours was not affected by ACTH infusion, consistent with overall sodium balance (**Figure 1F**). Sodium excretion has diurnal variation (**Figure 1G, left**) and physiologically ∼75% of the total output is achieved during the period of activity. ACTH disrupted this diurnal variation, with only ∼50% of the sodium output achieved during the active period (**Figure 1G, right**). The normal circadian variation in urine and potassium output was unaffected by ACTH treatment (**Supplemental figure 3**).

### ACTH impairs renal autoregulation and renal artery contractile responses

In separate groups of ACTH-treated and control mice, we used a Doppler flow probe to measure blood flow in the left renal artery at baseline and after increased BP. At baseline, renal blood flow was higher (**Figure 2A**) and renal vascular resistance lower (**Figure 2B**) in ACTH-treated mice. BP was then increased by arterial ligation, using the pressure-natriuresis protocol, and blood flow was measured throughout (**Figures 2C & 2D**). As BP increased, control mice autoregulated renal blood flow (**Figure 2E**) and flow was no different at peak BP compared with baseline (**Figure 2F**). Autoregulation was impaired in ACTH-infused mice and at peak BP, blood flow was significantly higher than at baseline. The autoregulatory index was significantly higher in ACTH-treated mice, indicative of reduced autoregulatory efficiency (**Figure 2G**). GFR again autoregulated effectively in both groups (**Figure 2H**), suggesting that hemodynamic impairment was not evident in the afferent arteriole, and was restricted to the larger renal arteries.

**Figure 2:**
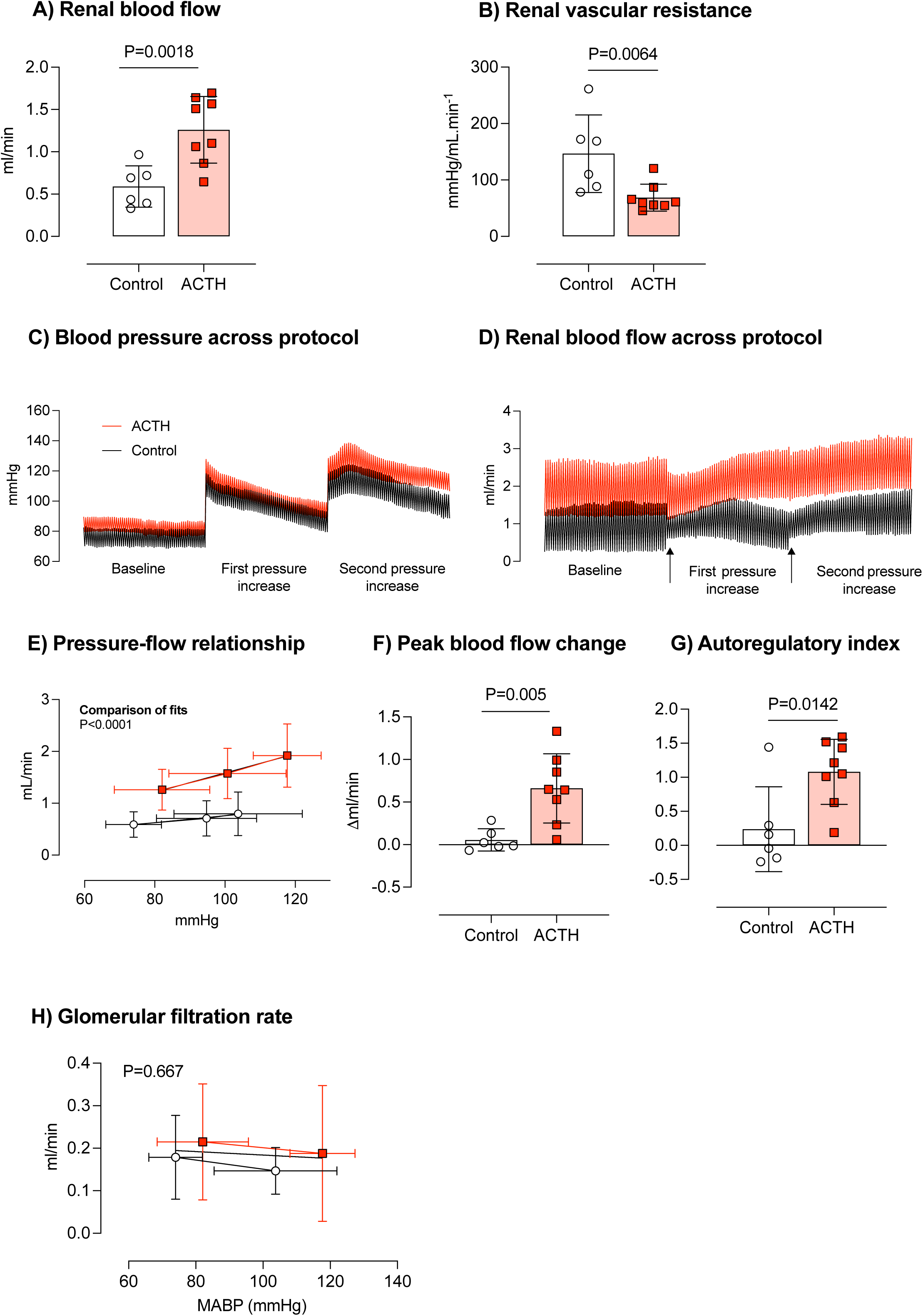
Renal hemodynamics. Mice were infused with ACTH (red square; n=8) or saline vehicle (controls; open circle; n=6) for 21 days and were then anesthetised to allow placement of a Doppler flow probe around the left renal artery at baseline and following two consecutive periods in which BP was acutely increased by arterial ligation. **A)** baseline renal blood flow and **B)** renal vascular resistance. Individual data points and group mean±SD are shown and statistical comparisons were made by unpaired t-test, with two-tailed P values given. **C)** An example of the BP profile over the duration of the experiment (left); and (right) the between BP and renal blood flow, shown as group mean±SD with liner regression curve relationship with P-value for the comparison of fits. **D)** the net change in renal blood flow, calculated by subtracting baseline values from the value recorded in the final period of the protocol. Individual data points with group mean±SD are shown and compared by unpaired t-test, with two-tailed P-value given. **E)** The relationship between BP and GFR, shown as group mean±SD.

To investigate this further, wire myography was used to examine *ex vivo* constriction and relaxation responses of the renal artery. The contractile response to the depolarization stimulus (125 mM extracellular potassium) was no different between control and ACTH-treated groups and did not deteriorate over the time course of the myography protocol. Normalized to this potassium response, renal arteries from ACTH-treated mice showed an attenuated constriction response to the adrenoceptor agonist, phenylephrine (**Figure 3A**). The maximal response (%Emax, Control: 142±2% *vs* ACTH:113±12%; *P*=0.0307) and the sensitivity (logEC_50_, Control: 6.66±0.04 *vs* ACTH: 5.81±0.21; *P*=0.0009), were both significantly reduced. Vasodilatory responses were also examined. The response to the endothelium-dependent vasodilator, acetylcholine, was impaired (**Figure 3B**), with significant reductions in both the sensitivity (logIC_50_, Control: 7.27±0.14 *vs* ACTH: 6.55±0.33; *P*=0.0009) and maximum vasodilation (Control: 5.42±3.60% *vs* ACTH: 42.41±7.25%; *P*<0.0001). The endothelium-independent dilation response to the nitric oxide donor, sodium nitroprusside was similarly impaired (**Figure 3C**), with significant reductions in sensitivity (logIC_50_, Control: 7.78±0.10 *vs* ACTH: 6.55±0.33; *P*=0.001) and maximum vasodilation (Control: -1.09±2.28% *vs* ACTH: 29.9±7.23%; *P*=0.0007).

**Figure 3:**
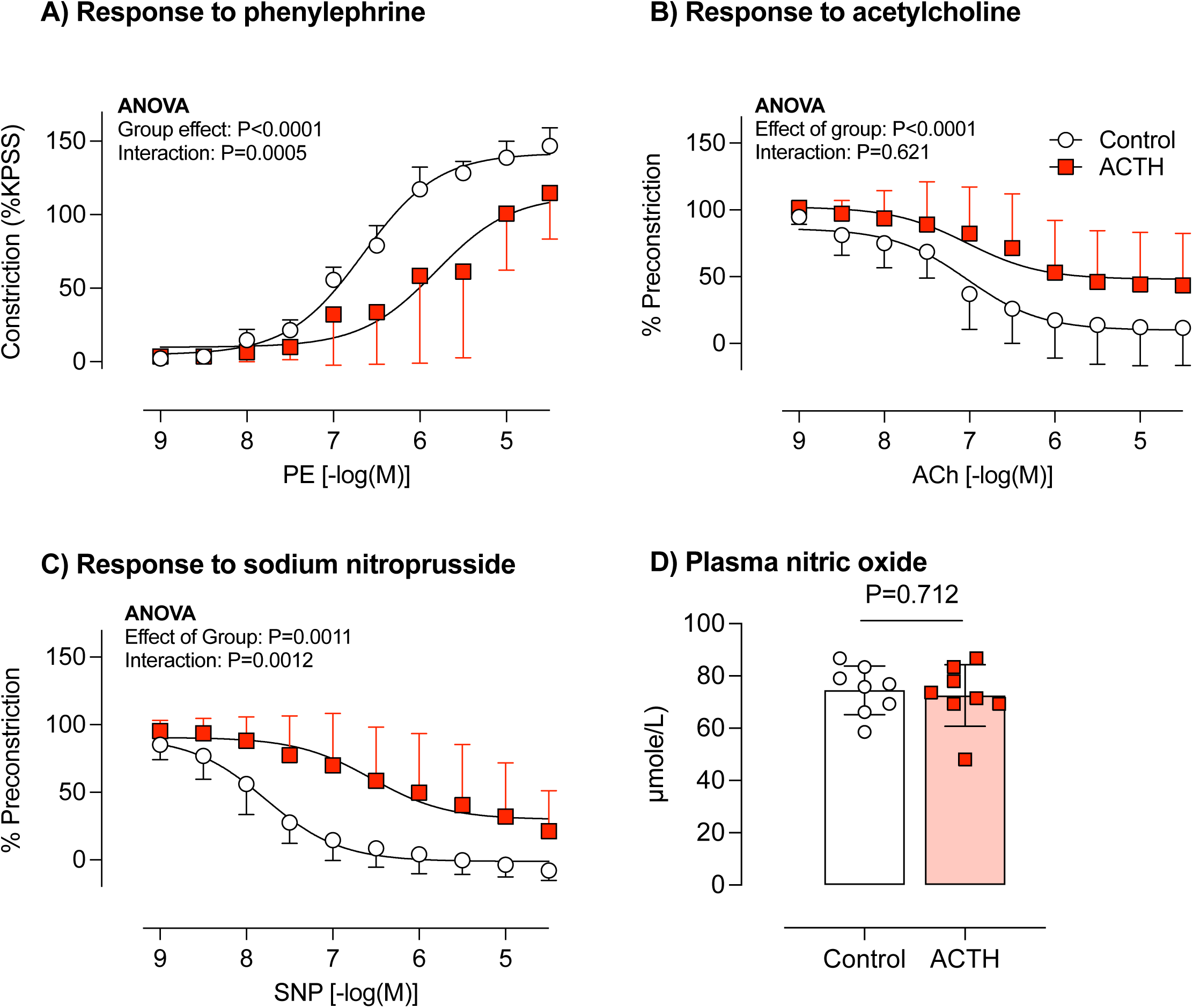
Ex vivo vascular function. Mice were infused with ACTH (red square; n=8) or saline vehicle (controls; open circle; n=6) for 14 day, killed between 7-8pm by decapitation, allowing rapid blood collection and isolation of the left renal artery. Artery rings were mounted on a wire myograph for measurement of **A)** the contractile response to phenylephrine, expressed as a % of the maximal constriction induced by 125 mmol/l KCL; the dilation induced by **B)** acetylcholine (endothelium-dependent) and **C)** sodium nitroprusside (endothelium-independent). Data are group mean ±SD and statistical comparisons were made by two-way ANOVA for the main effects of concentration, treatment group and interaction; P-values as shown. **D)** Plasma nitric oxide concentration. Individual data points are shown with group mean±SD and compared by unpaired t-test with two-tailed P-value as shown.

To explore these functional changes at a molecular level, we used qPCR to assess mRNA abundance for key transcripts in the renal artery (**Supplemental figure 4**). ACTH treatment did not affect mRNA abundance of *Adra1a* (encoding α1A adrenoreceptor), *Nos3* (endothelial nitric oxide synthase) or *Prkg1* (encoding α- and β-isoforms of soluble cyclic GMP-dependent protein kinase 1). *Nr3c1 (*GR) expression was downregulated in ACTH mice but *hsd11b1,* encoding the glucocorticoid regenerating enzyme 11βHSD1 was significantly increased: the upregulation of *fkbp5* mRNA (FK506 binding protein 5) is indicative of enhanced GR? signalling. Expression of *Nr3c2,* (MR) was not changed but *hsd11b2* was increased. We found evidence of remodelling, with downregulation of *Eln* (elastin) and *Emilin-1 (*EMILIN-1), alongside upregulation of *Vegfr2* (VEGFR2) and *Tgfb1* (TGFβ).

As a comparator for renal artery changes, we made these functional and molecular measurements in isolated mesenteric arteries. In this context, ACTH treatment did not change the contractile response to phenylephrine, acetylcholine or to sodium nitroprusside (**Supplemental figure 5**). *Fkbp5* mRNA abundance was significantly higher in ACTH-treated mice, but the other transcripts were unchanged (**Supplemental figure 6).**

### ACTH increases BP and induces non-dipping BP

We used radiotelemetry to measure the impact of ACTH on BP. Recordings were made in n=4 mice for three consecutive baseline days and over 21 days of ACTH infusion (**Figure 4A**). ACTH infusion increased BP and at the end of the protocol both 24h systolic BP (+7.0±1.5 mmHg; *P*<0.001; **Figure 4B**) and 24h diastolic BP (+2.1±0.6 mmHg; *P*=0.004; **Figure 4C**) were increased compared to baseline. At this point, ACTH infusion had also diminished the diurnal BP variation, attenuating the systolic and diastolic BP dip across the sleep phase of the 24h cycle (**Figure 4D**). This effect on BP variability did not reflect changes to locomotor activity (**Figure 4E**), which continued to dip in the sleep phase and was not affected by ACTH infusion (**Figure 4F**).

**Figure 4:**
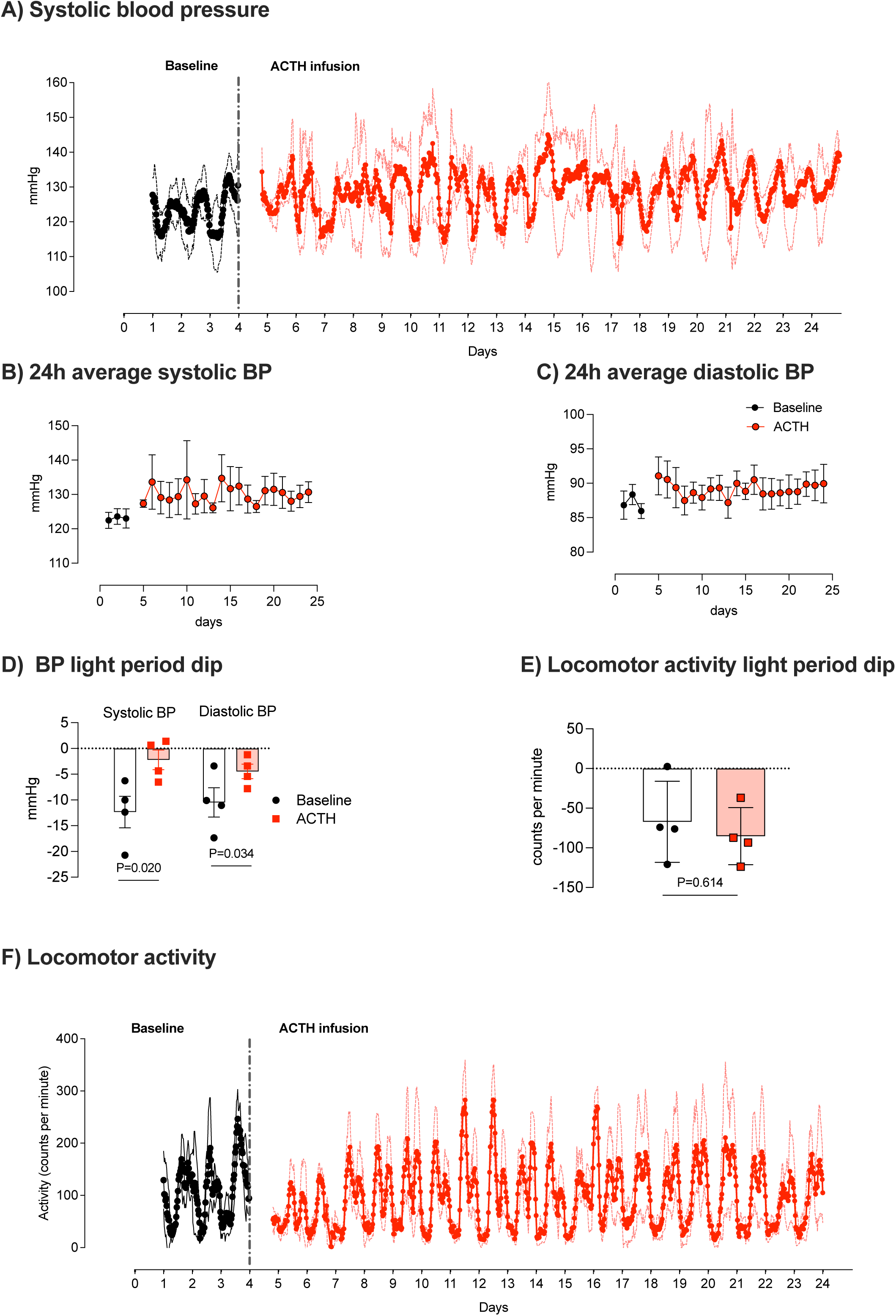
Blood pressure. Radiotelemetry was used to measure BP in conscious, unrestrained mice (n=4) for a 3-day baseline period (black circles) and for 21 days following infusion of ACTH (red circles). **A)** Systolic BP profile through the protocol, shown as group mean±SD. 24-hour average **B)** systolic BP and **C)** diastolic BP, presented as group mean±SD. **D)** the average systolic and diastolic BP dip during the light phase period; **E)** the average locomotor activity dip during the light phase period. Individual data points with group mean±SD are shown and compared by paired t-test, with two-tailed P-value given. **F)** Locomotor activity profile through the protocol, shown as group mean±SD.

### ACTH increases salt preference and amplifies salt-sensitive BP

ACTH infusion caused an array of phenotypes contributing to abnormal renal sodium excretion and BP and we next hypothesized that Cushing Syndrome mice would show an exaggerated BP response to high salt intake. Since ACTH can induce salt appetite, we first examined self-directed saline (1.5%) intake in mice at baseline and again after 21 days of infusion with either ACTH or vehicle. ACTH-treated mice showed increased intake of both saline and water (**Supplemental figure 7**).

In another group of mice (n=6), we used 3% Na food to give a matched high salt challenge, measuring the BP response before and after ACTH-infusion (**Figure 5A**). At baseline, the 24-hour average systolic BP was 122.7±2.4 mmHg and diastolic BP was 92.2±2.7 mmHg (**Figure 5B**). High salt intake significantly increased both systolic (+10.1±1.0 mmHg; *P*<0.001) and diastolic BP (+8.0±1.1 mmHg; *P*<0.001), without affecting the BP rhythmicity or dip during the light phase period of inactivity.

**Figure 5:**
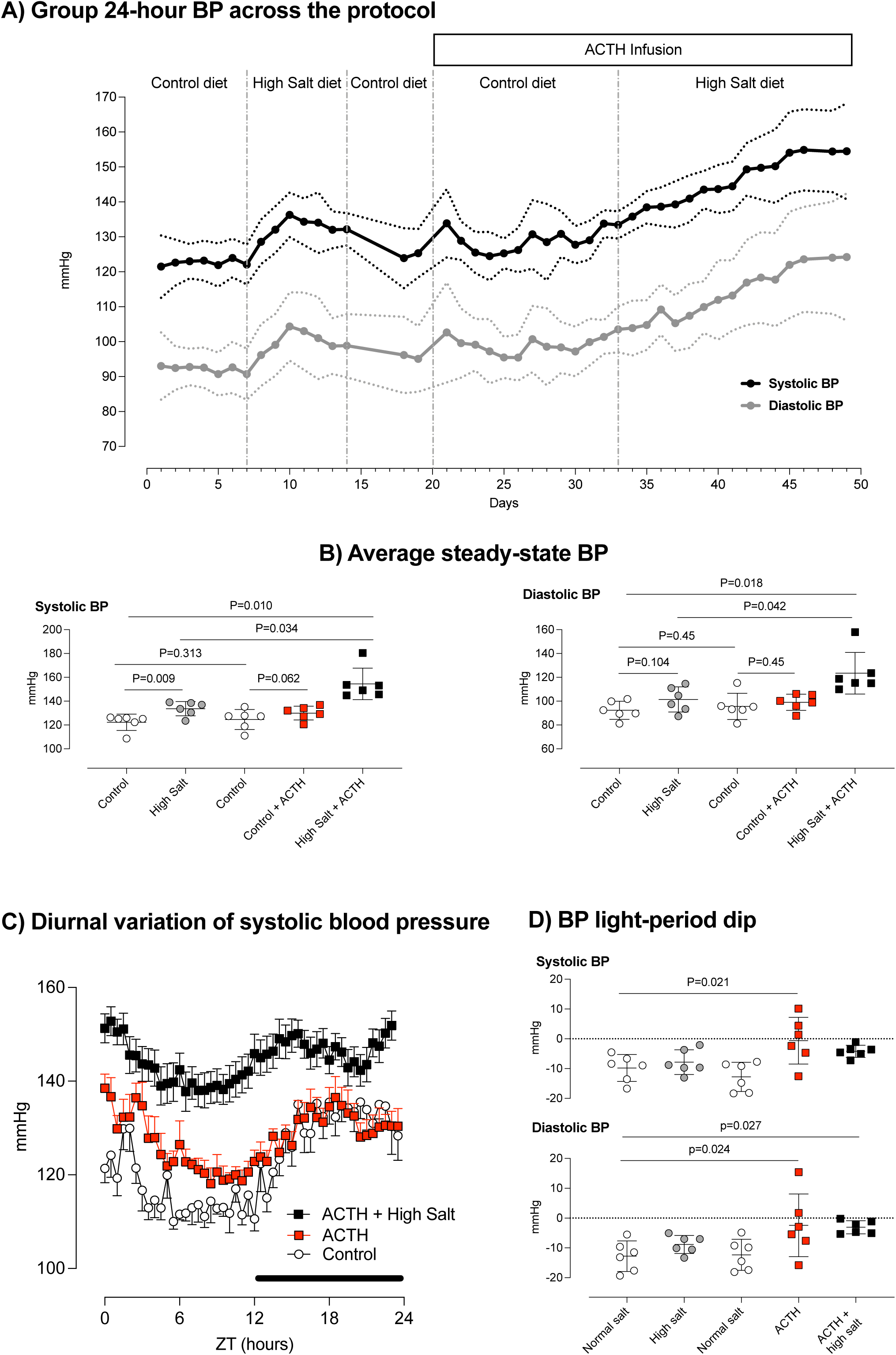
Salt-sensitivity of blood pressure. Radiotelemetry was used to measure BP in conscious, unrestrained mice (n=6) across a 50-day protocol to examine the effect of high salt intake on BP. **A)** systolic and diastolic BP through the treatment phases of the protocol, shown as group mean±SD. **B)** average steady-state systolic (left) and diastolic (right) BP. Individual data points are shown with group mean±SD and compared by ANOVA with Holm-Šidák for planned comparisons; P-values as shown. **C)** 24-hour BP profile during the control period (open circles), following ACTH infusion (red squares) and during the ACTH and high salt intake phase. **D)** the average systolic (top panel) and diastolic (bottom panel) BP dip during the light phase period; group mean±SD are shown and compared by ANOVA, with Holm-Šidák for planned comparisons.

Mice were returned to regular chow for a washout period during which BP returned to baseline values. ACTH was then given *via* osmotic minipump. In this group of mice there was no significant increase in 24-hour average systolic or diastolic BP, however, the time-of-day rhythmicity was dampened (**Figure 5C**), with loss of the BP dip (**Figure 5D**). In the final phase, ACTH infusion was maintained, and mice were returned to the high salt diet. ACTH infusion amplified salt-sensitivity, increasing systolic BP by 18.6±2.2 mmHg (*P*<0.001) and diastolic BP by 14.7±2.3 mmHg (*P*<0.001). Plasma steroid hormones were also measured (**Supplemental figure 8**). ACTH infusion increased plasma corticosterone, reversing the diurnal cycle; this was amplified by high salt intake. ACTH also increased circulating aldosterone and abolished the normal suppression of aldosterone by high salt intake.

## DISCUSSION

The main finding of our study in a mouse model of Cushing Syndrome was a substantially diminished pressure-natriuresis due to renal vascular and tubular abnormalities. On a normal salt intake, Cushing Syndrome impaired the diurnal rhythm of sodium excretion and BP. The capacity of high salt intake to suppress aldosterone was lost and ACTH-excess promoted a transition to salt-sensitive hypertension, which in humans increases cardiovascular and mortality risk^19,24^.

Pressure natriuresis uses intrarenal paracrine signalling to modulate sodium excretion in response to renal artery perfusion pressure. Blood flow in the renal medulla is key to pressure-natriuresis^21^ and an imbalance in any component diminishes its effectiveness^25,26^. It is a gateway to salt-sensitive hypertension, although whether this is through sodium retention^27^ or as a hallmark of generalised vasodysfunction^28^ is controversial. In our mouse model of Cushing Syndrome, the peak increase in fractional sodium excretion, an index of tubular reabsorption, was about half that of control animals. This is a significant failure to downregulate epithelial sodium transport. Our study does not identify the specific transporter(s) responsible but it is unlikely to be a single point of failure: glucocorticoid excess will activate GR^9,29,11^ and MR^14,17^ to promote sodium transport all along the nephron and can stimulate NCC and NKCC2 indirectly because of hypokalemia^15^. It is also possible that the primary abnormality lies within the vascular, or pressure-sensing part of the response, rather than the effector systems that lead to natriuresis. We found that glucocorticoid excess increased renal blood flow and reduced the ability to autoregulate blood flow against a rise in renal perfusion pressure. This is important for two reasons: first, barotrauma to glomeruli contributes to progressive renal injury^30^. Second, an abnormal autoregulatory index can signal a generalized hemodynamic dysfunction that attenuates the ability of the kidney to “sense” pressure. Dahl Salt-Sensitive rats, for example, also have an impaired autoregulatory index and renal blood flow rises with BP across the physiological range of BP. It might be anticipated that this would cause larger blood flows through the renal medulla and augment pressure natriuresis. However, the relationship is reset so that at any given BP, renal blood flow is higher and medullary flow is lower than in salt-resistant animals: the pressure natriuresis response is impaired^31,32^.

Our study did not measure medullary blood flow directly and because we did not use manoeuvres to reduce renal perfusion pressure below baseline values, the full autoregulatory range was not assessed. Nevertheless, our results point to abnormalities in myogenic tone which contribute to progressive autoregulatory failure^33^. Generalised vasodysfunction contributes importantly to salt-sensitivity^28^ and we therefore used isolated renal and mesenteric arteries to gain mechanistic insight into general vascular pathology. As anticipated, there was a molecular signature of increased glucocorticoid exposure in both arteries but only the renal artery showed an attenuated vasodilation response to acetylcholine. Notably, Cushing patients can show reduced nitric oxide bioavailability and animal models of glucocorticoid excess provide evidence of endothelial dysfunction^4^. However, this is not evident in our model and the impaired dilation response to the nitric oxide donor instead point to a vascular smooth muscle cell defect. In this context, conditional deletion of GR in vascular smooth muscle protects against dexamethasone induced hypertension.^34^ It is possible that our model captures early vascular pathophysiology and that the conduit renal artery is more susceptible to this than the mesenteric resistance arteries. In this context, we assessed mRNA for VEGF receptor 2, because VEGF is elevated along with other markers of arterial remodelling in Cushing patients, more so than in hypertension or Conns disease^35^. *Vegfr2* was increased only in the renal artery; reduced elastin and increased TGFB mRNA provide other hallmarks of remodelling. An interesting observation was reduced expression of Emilin-1, a structural protein of the elastic fibre expressed in the adventitia. Emilin-1 acts as an endogenous brake on TGFB signalling and loss of function disrupts the collagen network reducing vascular elastogenesis^36^ and, myogenic tone^37^: knockout mice are hypertensive^38^.

Impaired pressure-natriuresis response is proposed as a core causal mechanism for the development of salt-sensitive hypertension, driving sodium retention and ECFV expansion^27^. Consistent with this are studies in healthy humans in which chronic infusion of either ACTH or cortisol causes sodium retention and volume expansion^16^. However, our studies were performed after sustained ACTH excess and we have no evidence at this time point for ECFV expansion. Indeed, the mice had hypernatremia, plasma hypertonicity and elevated copeptin, all of which are indicative of ECFV contraction rather than volume expansion. However, even with the low equivalent sodium intake provided by regular rodent diets, our studies identify a reduced capacity to rapidly excrete salt. The diurnal pattern of sodium excretion was disrupted by ACTH excess and relatively more sodium was excreted during the sleep phase. Given that glucocorticoids are important circadian hormones^39^, it is possible that a disrupted sleep/eat cycle contributes to the changed sodium excretion pattern. However, two things argue against this. First, telemetry recording of activity and heart rate indicate that the normal diurnal variation of the sleep/wake cycle was maintained during ACTH infusion; second, urine flow and potassium excretion retained normal diurnal variation, suggesting that the circadian abnormality is specific for salt.

The increased excretion of sodium during the sleep phase was parallelled by disrupted BP rhythmicity, with attenuation of the “dip”. Non-dipping BP is an independent cardiovascular risk factor^40^ and commonly observed in Cushing patients, particularly those with an adrenal origin for the disease^4^. These circadian abnormalities in salt excretion and BP are seen in human studies and have been attributed to an impaired pressure natriuresis response: a diminished capacity to efficiently excrete salt during the day causes BP to remain elevated at night to excrete the residual salt and achieve balance^41–43^.

Our model of Cushing Syndrome displayed renal and cardiovascular abnormalities suggesting loss of resilience in salt homeostasis and we hypothesized that the mice would show a strong salt sensitive BP. Examining the response to a high salt challenge in a single group of mice before and after chronic ACTH infusion revealed that C57BL6 mice, which are often considered a salt-resistant strain^44^, have a degree of salt-sensitivity. Others have reported this^45^ and our previous work indicates that the salt-sensitivity reflects increased sympathetic drive and not impaired pressure natriuresis and salt retention^46^. ACTH infusion amplified the BP response to high salt diet, supporting our hypothesis. Furthermore, despite persistent hypokalemia, ACTH-infusion abolished the suppression of aldosterone by salt intake. This might reflect a direct stimulatory effect of ACTH on aldosterone synthase^47^, overriding the normal drivers of aldosterone regulation.

In summary, the mouse model of Cushing Syndrome displays reduced ability of the kidney to regulate BP *via* salt excretion. Activated salt preference – increased salt intake in the salt replete state – may reflect activation of central pathways linked to gratification of primal needs^48^ and would potentiate the vulnerability to salt loads. There are further confounding factors: a single high salt meal stimulates ACTH and cortisol production^1^ and chronic high salt intake modulates CNS and peripheral pathways to enhance circulating and tissue glucocorticoid bio-availability^49^.

Given that high salt intake is the societal norm worldwide^50^, this intersection of detrimental salt homeostasis phenotypes can create a vicious cycle impacting BP. Hypertension accounts for much of the cardiovascular disease burden in Cushing Syndrome^1,3,4^. Salt-sensitivity^19^ and non-dipping BP^40,51^ are interrelated morbidities that add to this adverse risk profile.

## Acknowledgements and Funding Sources

Research funding for this study came from The British Heart Foundation (FS/16/54/32730; PG/16/98/32568; RE/18/5/34216); Kidney Research UK (IN001/2017; INT001/2018) and a Chief Scientist Office Senior Clinical Research Fellowship (SCAF/19/02). For the purpose of open access, the author has applied a Creative Commons Attribution (CC BY) licence to any Author Accepted Manuscript version arising from this submission.

## Author Contributions

HMC, JRI, MCH, DEWL, ND and MAB contributed to the study design. HMC, JRI, CG, KS designed protocols and performed experiments. All authors contributed to data production, statistical analysis, and interpretation. MAB and ND wrote the initial draft of the manuscript; all authors reviewed and revised the manuscript.

## REFERENCES

1. Pivonello R, Isidori AM, De Martino MC, Newell-Price J, Biller BM, Colao A. Complications of Cushing’s syndrome: state of the art. Lancet Diabetes Endocrinol. 2016;4:611–629. doi: 10.1016/S2213-8587(16)00086-3

2. Menzies-Gow AN, Tran TN, Stanley B, Carter VA, Smolen JS, Bourdin A, Fitzgerald JM, Raine T, Chapaneri J, Emmanuel B, et al. Trends in Systemic Glucocorticoid Utilization in the United Kingdom from 1990 to 2019: A Population-Based, Serial Cross-Sectional Analysis. Pragmat Obs Res. 2024;15:53–64. doi: 10.2147/POR.S442959

3. Fardet L, Petersen I, Nazareth I. Risk of cardiovascular events in people prescribed glucocorticoids with iatrogenic Cushing’s syndrome: cohort study. BMJ. 2012;345:e4928. doi: 10.1136/bmj.e4928

4. Isidori AM, Graziadio C, Paragliola RM, Cozzolino A, Ambrogio AG, Colao A, Corsello SM, Pivonello R, Group ABCS. The hypertension of Cushing’s syndrome: controversies in the pathophysiology and focus on cardiovascular complications. J Hypertens. 2015;33:44–60. doi: 10.1097/HJH.0000000000000415

5. Hunter RW, Ivy JR, Bailey MA. Glucocorticoids and renal Na+ transport: implications for hypertension and salt sensitivity. J Physiol. 2014;592:1731–1744. doi: 10.1113/jphysiol.2013.267609

6. Yasuda G, Shionoiri H, Umemura S, Takasaki I, Ishii M. Exaggerated blood pressure response to angiotensin II in patients with Cushing’s syndrome due to adrenocortical adenoma. Eur J Endocrinol. 1994;131:582–588. doi: 10.1530/eje.0.1310582

7. Heaney AP, Hunter SJ, Sheridan B, Brew Atkinson A. Increased pressor response to noradrenaline in pituitary dependent Cushing’s syndrome. Clin Endocrinol (Oxf). 1999;51:293–299. doi: 10.1046/j.1365-2265.1999.00766.x

8. Bailey MA. 11beta-Hydroxysteroid Dehydrogenases and Hypertension in the Metabolic Syndrome. Curr Hypertens Rep. 2017;19:100. doi: 10.1007/s11906-0170797-z

9. Bobulescu IA, Dwarakanath V, Zou L, Zhang J, Baum M, Moe OW. Glucocorticoids acutely increase cell surface Na+/H+ exchanger-3 (NHE3) by activation of NHE3 exocytosis. Am J Physiol Renal Physiol. 2005;289:F685–691. doi: 10.1152/ajprenal.00447.2004

10. Attmane-Elakeb A, Sibella V, Vernimmen C, Belenfant X, Hebert SC, Bichara M. Regulation by glucocorticoids of expression and activity of rBSC1, the Na+-K+(NH4+)-2Cl-cotransporter of medullary thick ascending limb. J Biol Chem. 2000;275:33548–33553. doi: 10.1074/jbc.M006591200

11. Ivy JR, Jones NK, Costello HM, Mansley MK, Peltz TS, Flatman PW, Bailey MA. Glucocorticoid receptor activation stimulates the sodium-chloride cotransporter and influences the diurnal rhythm of its phosphorylation. Am J Physiol Renal Physiol. 2019;317:F1536–F1548. doi: 10.1152/ajprenal.00372.2019

12. Ackermann D, Gresko N, Carrel M, Loffing-Cueni D, Habermehl D, Gomez-Sanchez C, Rossier BC, Loffing J. In vivo nuclear translocation of mineralocorticoid and glucocorticoid receptors in rat kidney: differential effect of corticosteroids along the distal tubule. Am J Physiol Renal Physiol. 2010;299:F1473–1485. doi: 10.1152/ajprenal.00437.2010

13. Gaeggeler HP, Gonzalez-Rodriguez E, Jaeger NF, Loffing-Cueni D, Norregaard R, Loffing J, Horisberger JD, Rossier BC. Mineralocorticoid versus glucocorticoid receptor occupancy mediating aldosterone-stimulated sodium transport in a novel renal cell line. J Am Soc Nephrol. 2005;16:878–891. doi: 10.1681/ASN.2004121110

14. Loughlin S, Costello HM, Roe AJ, Buckley C, Wilson SM, Bailey MA, Mansley MK. Mapping the Transcriptome Underpinning Acute Corticosteroid Action within the Cortical Collecting Duct. Kidney360. 2023;4:226–240. doi: 10.34067/KID.0003582022

15. Salih M, Bovee DM, van der Lubbe N, Danser AHJ, Zietse R, Feelders RA, Hoorn EJ. Increased Urinary Extracellular Vesicle Sodium Transporters in Cushing Syndrome With Hypertension. J Clin Endocrinol Metab. 2018;103:2583–2591. doi: 10.1210/jc.2018-00065

16. Connell JM, Whitworth JA, Davies DL, Lever AF, Richards AM, Fraser R. Effects of ACTH and cortisol administration on blood pressure, electrolyte metabolism, atrial natriuretic peptide and renal function in normal man. J Hypertens. 1987;5:425–433.

17. Bailey MA, Mullins JJ, Kenyon CJ. Mineralocorticoid and glucocorticoid receptors stimulate epithelial sodium channel activity in a mouse model of Cushing syndrome. Hypertension. 2009;54:890–896. doi: 10.1161/HYPERTENSIONAHA.109.134973

18. Dunbar DR, Khaled H, Evans LC, Al-Dujaili EA, Mullins LJ, Mullins JJ, Kenyon CJ, Bailey MA. Transcriptional and physiological responses to chronic ACTH treatment by the mouse kidney. Physiol Genomics. 2010;40:158–166. doi: 10.1152/physiolgenomics.00088.2009

19. Bailey MA, Dhaun N. Salt Sensitivity: Causes, Consequences, and Recent Advances. Hypertension. 2024;81:476–489. doi: 10.1161/HYPERTENSIONAHA.123.17959

20. Ivy JR, Bailey MA. Pressure natriuresis and the renal control of arterial blood pressure. J Physiol. 2014;592:3955–3967. doi: 10.1113/jphysiol.2014.271676

21. Cowley AW, Jr., Roman RJ, Mattson DL, Franchini KG, O’Connor PM, Makino A, Taylor NE, Evans LC, Mori T, Dickhout JG, et al. Renal Medulla in Hypertension. Hypertension. 2024;81:2383–2394. doi: 10.1161/HYPERTENSIONAHA.124.21711

22. Menzies RI, Zhao X, Mullins LJ, Mullins JJ, Cairns C, Wrobel N, Dunbar DR, Bailey MA, Kenyon CJ. Transcription controls growth, cell kinetics and cholesterol supply to sustain ACTH responses. Endocr Connect. 2017;6:446–457. doi: 10.1530/EC-17-0092

23. Al-Dujaili EA, Mullins LJ, Bailey MA, Andrew R, Kenyon CJ. Physiological and pathophysiological applications of sensitive ELISA methods for urinary deoxycorticosterone and corticosterone in rodents. Steroids. 2009;74:938–944. doi: 10.1016/j.steroids.2009.06.009

24. Nishimoto M, Griffin KA, Wynne BM, Fujita T. Salt-Sensitive Hypertension and the Kidney. Hypertension. 2024;81:1206–1217. doi: 10.1161/HYPERTENSIONAHA.123.21369

25. Culshaw GJ, Costello HM, Binnie D, Stewart KR, Czopek A, Dhaun N, Hadoke PWF, Webb DJ, Bailey MA. Impaired pressure natriuresis and non-dipping blood pressure in rats with early type 1 diabetes mellitus. J Physiol. 2019;597:767–780. doi: 10.1113/JP277332

26. Sipos A, Vargas SL, Toma I, Hanner F, Willecke K, Peti-Peterdi J. Connexin 30 deficiency impairs renal tubular ATP release and pressure natriuresis. J Am Soc Nephrol. 2009;20:1724–1732. doi: 10.1681/ASN.2008101099

27. Hall JE. Renal Dysfunction, Rather Than Nonrenal Vascular Dysfunction, Mediates Salt-Induced Hypertension. Circulation. 2016;133:894–906. doi: 10.1161/CIRCULATIONAHA.115.018526

28. Morris RC, Jr., Schmidlin O, Sebastian A, Tanaka M, Kurtz TW. Vasodysfunction That Involves Renal Vasodysfunction, Not Abnormally Increased Renal Retention of Sodium, Accounts for the Initiation of Salt-Induced Hypertension. Circulation. 2016;133:881–893. doi: 10.1161/CIRCULATIONAHA.115.017923

29. Loffing J, Lotscher M, Kaissling B, Biber J, Murer H, Seikaly M, Alpern RJ, Levi M, Baum M, Moe OW. Renal Na/H exchanger NHE-3 and Na-PO4 cotransporter NaPi-2 protein expression in glucocorticoid excess and deficient states. J Am Soc Nephrol. 1998;9:1560–1567. doi: 10.1681/ASN.V991560

30. Mori T, Polichnowski A, Glocka P, Kaldunski M, Ohsaki Y, Liang M, Cowley AW, Jr. High perfusion pressure accelerates renal injury in salt-sensitive hypertension. J Am Soc Nephrol. 2008;19:1472–1482. doi: 10.1681/ASN.2007121271

31. Roman RJ, Kaldunski M. Pressure natriuresis and cortical and papillary blood flow in inbred Dahl rats. Am J Physiol. 1991;261:R595–602. doi: 10.1152/ajpregu.1991.261.3.R595

32. Roman RJ. Abnormal renal hemodynamics and pressure-natriuresis relationship in Dahl salt-sensitive rats. Am J Physiol. 1986;251:F57–65. doi: 10.1152/ajprenal.1986.251.1.F57

33. Carlstrom M. Nitric oxide signalling in kidney regulation and cardiometabolic health. Nat Rev Nephrol. 2021;17:575–590. doi: 10.1038/s41581-021-00429-z

34. Goodwin JE, Zhang J, Geller DS. A critical role for vascular smooth muscle in acute glucocorticoid-induced hypertension. J Am Soc Nephrol. 2008;19:1291–1299. doi: 10.1681/ASN.2007080911

35. Zacharieva S, Atanassova I, Orbetzova M, Kirilov G, Nachev E, Kalinov K, Shigarminova R. Vascular endothelial growth factor (VEGF), prostaglandin E2(PGE2) and active renin in hypertension of adrenal origin. J Endocrinol Invest. 2004;27:742–746. doi: 10.1007/BF03347516

36. Adamo CS, Beyens A, Schiavinato A, Keene DR, Tufa SF, Morgelin M, Brinckmann J, Sasaki T, Niehoff A, Dreiner M, et al. EMILIN1 deficiency causes arterial tortuosity with osteopenia and connects impaired elastogenesis with defective collagen fibrillogenesis. Am J Hum Genet. 2022;109:2230–2252. doi: 10.1016/j.ajhg.2022.10.010

37. Carnevale D, Facchinello N, Iodice D, Bizzotto D, Perrotta M, De Stefani D, Pallante F, Carnevale L, Ricciardi F, Cifelli G, et al. Loss of EMILIN-1 Enhances Arteriolar Myogenic Tone Through TGF-beta (Transforming Growth Factor-beta)-Dependent Transactivation of EGFR (Epidermal Growth Factor Receptor) and Is Relevant for Hypertension in Mice and Humans. Arterioscler Thromb Vasc Biol. 2018;38:2484–2497. doi: 10.1161/ATVBAHA.118.311115

38. Litteri G, Carnevale D, D’Urso A, Cifelli G, Braghetta P, Damato A, Bizzotto D, Landolfi A, Ros FD, Sabatelli P, et al. Vascular smooth muscle Emilin-1 is a regulator of arteriolar myogenic response and blood pressure. Arterioscler Thromb Vasc Biol. 2012;32:2178–2184. doi: 10.1161/ATVBAHA.112.254664

39. Lightman SL, Conway-Campbell BL. Circadian and ultradian rhythms: Clinical implications. J Intern Med. 2024;296:121–138. doi: 10.1111/joim.13795

40. Ivy JR, Bailey MA. Nondipping Blood Pressure: Predictive or Reactive Failure of Renal Sodium Handling? Physiology (Bethesda). 2021;36:21–34. doi: 10.1152/physiol.00024.2020

41. Fukuda M, Mizuno M, Yamanaka T, Motokawa M, Shirasawa Y, Nishio T, Miyagi S, Yoshida A, Kimura G. Patients with renal dysfunction require a longer duration until blood pressure dips during the night. Hypertension. 2008;52:1155–1160. doi: 10.1161/HYPERTENSIONAHA.108.115329

42. Guerrot D, Hansel B, Perucca J, Roussel R, Bouby N, Girerd X, Bankir L. Reduced insulin secretion and nocturnal dipping of blood pressure are associated with a disturbed circadian pattern of urine excretion in metabolic syndrome. J Clin Endocrinol Metab. 2011;96:E929–933. doi: 10.1210/jc.2010-2337

43. Bankir L, Bochud M, Maillard M, Bovet P, Gabriel A, Burnier M. Nighttime blood pressure and nocturnal dipping are associated with daytime urinary sodium excretion in African subjects. Hypertension. 2008;51:891–898. doi: 10.1161/HYPERTENSIONAHA.107.105510

44. Lerman LO, Kurtz TW, Touyz RM, Ellison DH, Chade AR, Crowley SD, Mattson DL, Mullins JJ, Osborn J, Eirin A, et al. Animal Models of Hypertension: A Scientific Statement From the American Heart Association. Hypertension. 2019;73:e87–e120. doi: 10.1161/HYP.0000000000000090

45. Combe R, Mudgett J, El Fertak L, Champy MF, Ayme-Dietrich E, Petit-Demouliere B, Sorg T, Herault Y, Madwed JB, Monassier L. How Does Circadian Rhythm Impact Salt Sensitivity of Blood Pressure in Mice? A Study in Two Close C57Bl/6 Substrains. PLoS One. 2016;11:e0153472. doi: 10.1371/journal.pone.0153472

46. Ralph AF, Grenier C, Costello HM, Stewart K, Ivy JR, Dhaun N, Bailey MA. Activation of the Sympathetic Nervous System Promotes Blood Pressure Salt-Sensitivity in C57BL6/J Mice. Hypertension. 2021;77:158–168. doi: 10.1161/HYPERTENSIONAHA.120.16186

47. MacKenzie SM, van Kralingen JC, Davies E. Regulation of Aldosterone Secretion. Vitam Horm. 2019;109:241–263. doi: 10.1016/bs.vh.2018.07.001

48. Liedtke WB, McKinley MJ, Walker LL, Zhang H, Pfenning AR, Drago J, Hochendoner SJ, Hilton DL, Lawrence AJ, Denton DA. Relation of addiction genes to hypothalamic gene changes subserving genesis and gratification of a classic instinct, sodium appetite. Proc Natl Acad Sci U S A. 2011;108:12509–12514. doi: 10.1073/pnas.1109199108

49. Costello HM, Krilis G, Grenier C, Severs D, Czopek A, Ivy JR, Nixon M, Holmes MC, Livingstone DEW, Hoorn EJ, et al. High salt intake activates the hypothalamic-pituitary-adrenal axis, amplifies the stress response, and alters tissue glucocorticoid exposure in mice. Cardiovasc Res. 2023;119:1740–1750. doi: 10.1093/cvr/cvac160

50. Hunter RW, Dhaun N, Bailey MA. The impact of excessive salt intake on human health. Nat Rev Nephrol. 2022;18:321–335. doi: 10.1038/s41581-021-00533-0

51. Lempiainen PA, Ylitalo A, Huikuri H, Kesaniemi YA, Ukkola OH. Non-dipping blood pressure pattern is associated with cardiovascular events in a 21-year follow-up study. J Hum Hypertens. 2024;38:444–451. doi: 10.1038/s41371-024-00909-2

