## Supplemental Tables and Figures for "Renal arterial dysfunction, impaired pressure natriuresis and salt-sensitivity in a mouse model of Cushing syndrome"

**Table S1: Primers and probes for quantitative RT-PCR. Probe refers to the Universal Probe library (Roche UK)**

| Protein | Gene | Forward Primer | Reverse Primer | Probe |
|---|---|---|---|---|
| 11βHSD1 | *Hsd11b1* | tctacaaatgaagagttcagaccag | gccccagtgacaatcacttt | 1 |
| 11βHSD2 | *Hsd11b2* | cactcgaggggacgtattgt | gcaggggtatggcatgtct | 26 |
| 18S | *Rn18s* | gccgctagaggtgaaattctt | cgtcttcgaacctccgact | 93 |
| Adrenergic receptor alpha 1a | *Adra1a* | actgaaggtccgcttctcct | ggaatatttgctgagaccgaag | 1 |
| βactin | *Actb* | ctaaggccaaccgtgaaaag | accagaggcatacagggaca | 64 |
| Elastin | *Eln* | tgcagtactgtaaccccgttc | aggtggggtggtgagacat | 1 |
| Elastin microfibril interfacer 1 | *Emilin1* | cctgtctggctccagtgc | gctctagctgctgcaccttc | 46 |
| endothelial NOS | *Nos3* | atccagtgccctgcttca | gcagggcaagttaggatcag | 12 |
| FK506 binding protein 51 | *Fkbp5* | aaacgaaggagcaacggtaa | tcaaatgtccttccaccaca | 97 |
| GR | *Nr3c1* | gacgtgtggaagctgtaaagt | catttcttccagcacaaaggt | 56 |
| HPRT | *Hprt* | tcctcctcagaccgctttt | aacctggttcatcatcgctaa | 95 |
| MR | *Nr3c2* | ttcggagaaagaactgtcctg | cccagcttctttgactttcg | 50 |
| Protein kinase cGMP-dependent type 1 | *Prkg1* | acattccagagccttcctga | tctccattttcatagtgggtctc | 73 |
| TBP | *Tbp* | gggagaatcatggaccagaa | gatgggaattccaggagtca | 97 |
| TGF-β1 | *Tgf-b1* | ccctactctccccaagctgt | tgtgaagtggtggtctccag | 108 |

**Table S2**: Plasma parameters from male C57Bl6 mice treated with either ACTH (n=8) or vehicle control (n=9) for 21 days. Data are mean±SD. Statistical analysis was performed with unpaired T-test and the two-tailed P value is given.

| | Vehicle | ACTH | P-value |
|---|---|---|---|
| Sodium (mmol/l) | 147.2±3.1 | 149.5±4.4 | 0.312 |
| Potassium (mmol/l) | 4.17±0.32 | 3.70±0.51 | 0.015 |
| Osmolality (mOsm.kg) | 354±7 | 372±9 | 0.002 |
| Hematocrit (%) | 39.2±1.4 | 39.3±6 | 0.976 |
| Copeptin (pg/ml) | 15.8±10.6 | 92.3±30.6 | <0.0001 |

**Figure S1**. Mice were infused with ACTH (red square; n=7) or saline vehicle (controls; open circle; n=9) for 21 days. A) Body weight; B) weight of the gonadal fat pad; C) examples of adrenal gland hypertrophy; D) adrenal gland weight and cross sectional area; Plasma E) corticosterone and F) aldosterone. Individual data points and group mean±SD are shown. Statistical comparisons were made by unpaired t-test, with two-tailed P values shown

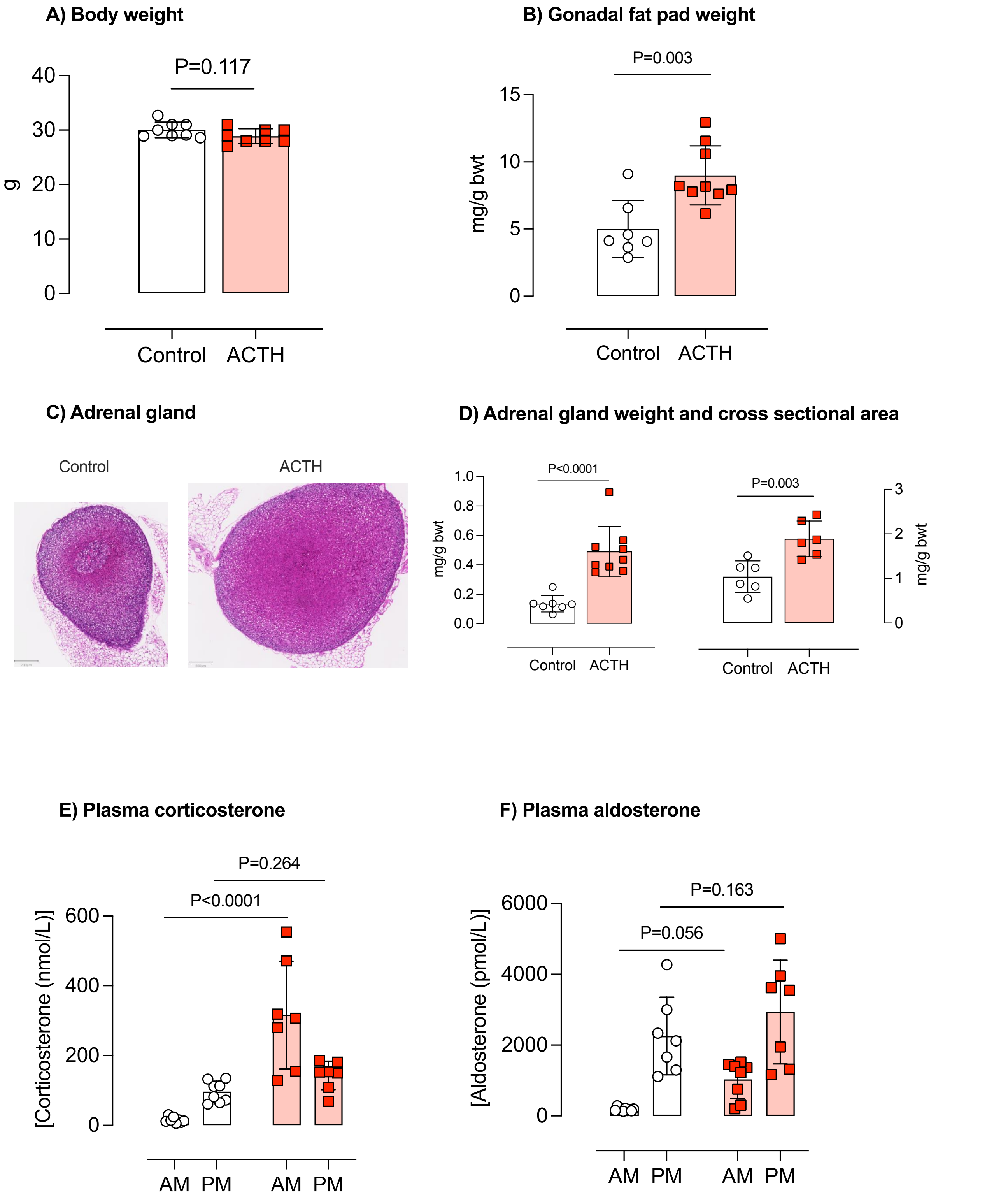

**Figure S2**. Mice were infused with ACTH (red square; n=7) or saline vehicle (controls; open circle; n=9) for 21 days and anaesthetised for measurement of A) pressure natriuresis; and B) pressure diuresis. Data are group mean±SD. Linear regression was used to compare curve fits between groups. C) The change in fractional sodium excretion was calculated by subtracting baseline values from the value obtained at peak pressure. Individual values are shown with group mean±SD. Fractional sodium excretion was significantly elevated over baseline in both groups (one sample t-test against "zero change"; P<0.0001); the net response was significantly greater in control compared with ACTH-treated mice., P value as shown.

**A) Pressure natriuresis response**

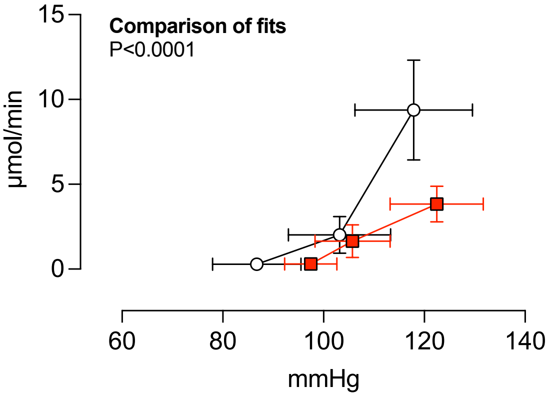

**B) Pressure diuresis response**

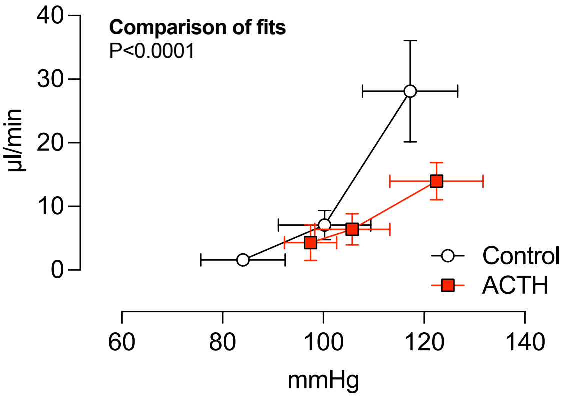

**C) Change in Fractional sodium excretion**

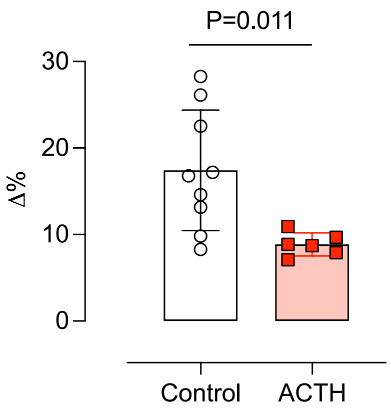

**Figure S3**. Mice (n=7) were housed individually in metabolism cages for measurement of food intake, water intake and urine output before (Control) and after (ACTH) 21-days of ACTH infusion. Urine was collected in two vials, one for the 12h period of darkness (active period) and 12h period of light (sleep period). Individual data points are shown along with group mean±SD. Statistical comparisons were made by paired-t-test, with P-values as shown.

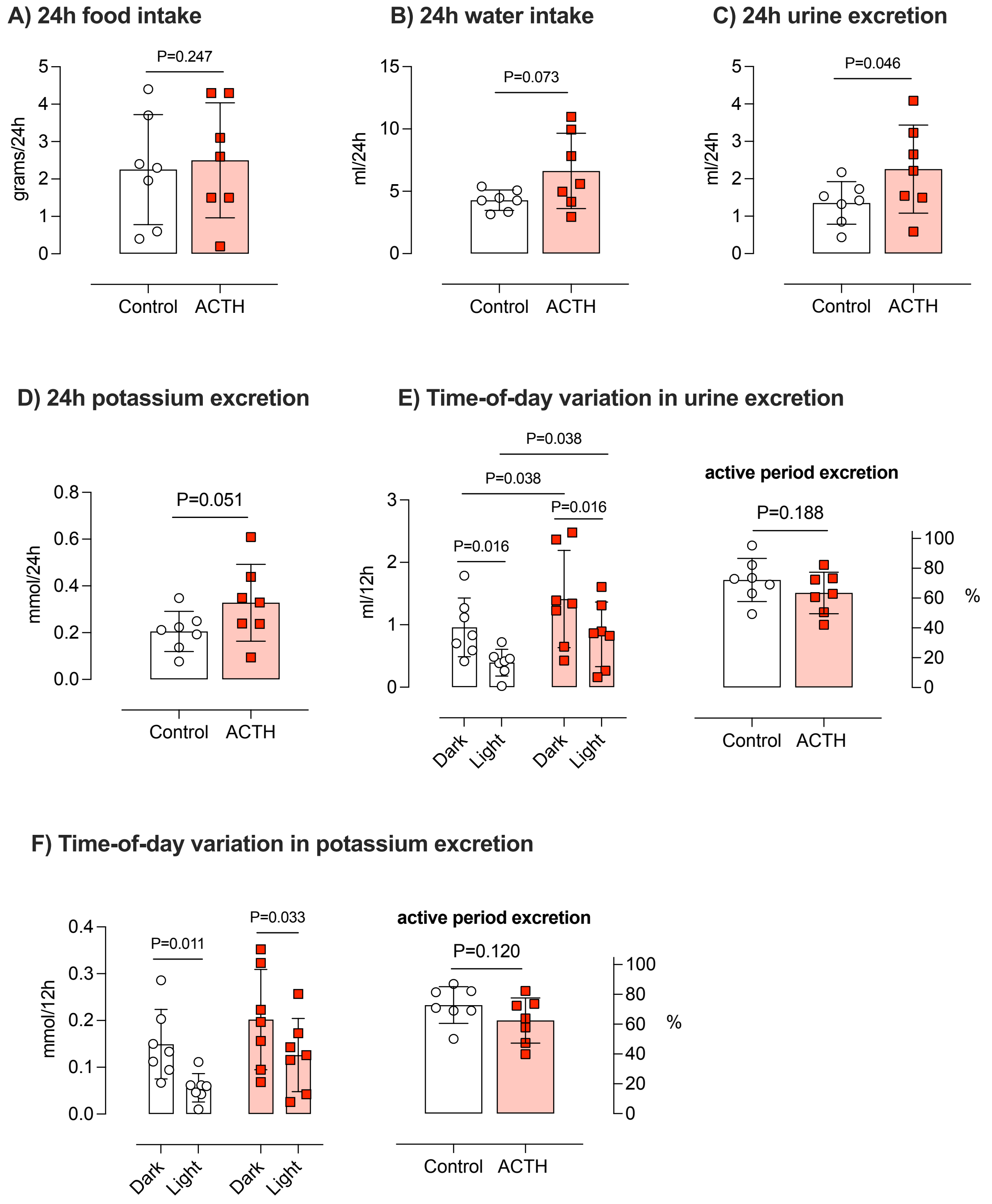

**Figure S4**. mRNA abundance of different genes in the renal artery isolated from mice treated for 14 days with vehicle control (n=4; open symbols) or ACTH (n=6; red circles). Target genes were *Adra1a* (encoding $\alpha$1A adrenoreceptor), *Nos3* (endothelial nitric oxide synthase), *Prkg1* (encoding $\alpha$- and $\beta$- isoforms of soluble cyclic GMP-dependent protein kinase 1), *Fkbp5* (FK506 binding protein 5), *Nr3c1* (glucocorticoid receptor), *Hsd11b1* (11$\beta$-hydroxysteroid dehydrogenase type 1), *Nr3c2* (mineralocorticoid receptor), *Hsd11b2* (11$\beta$-hydroxysteroid dehydrogenase type 2), *Eln* (elastin), *Emilin-1*, *Vegfr2* and *Tgf-b1.* A panel of appropriate reference (housekeeper; HK) genes, *Hprt*, *Rn18s*, *Actb* and *Tbp*, which did not differ between treatment groups. Data were log transformed to ensure symmetry of the data set and normality tests were conducted. Individual data points are shown, with group mean±SD. Comparisons were made with unpaired t-test; P-values as shown.

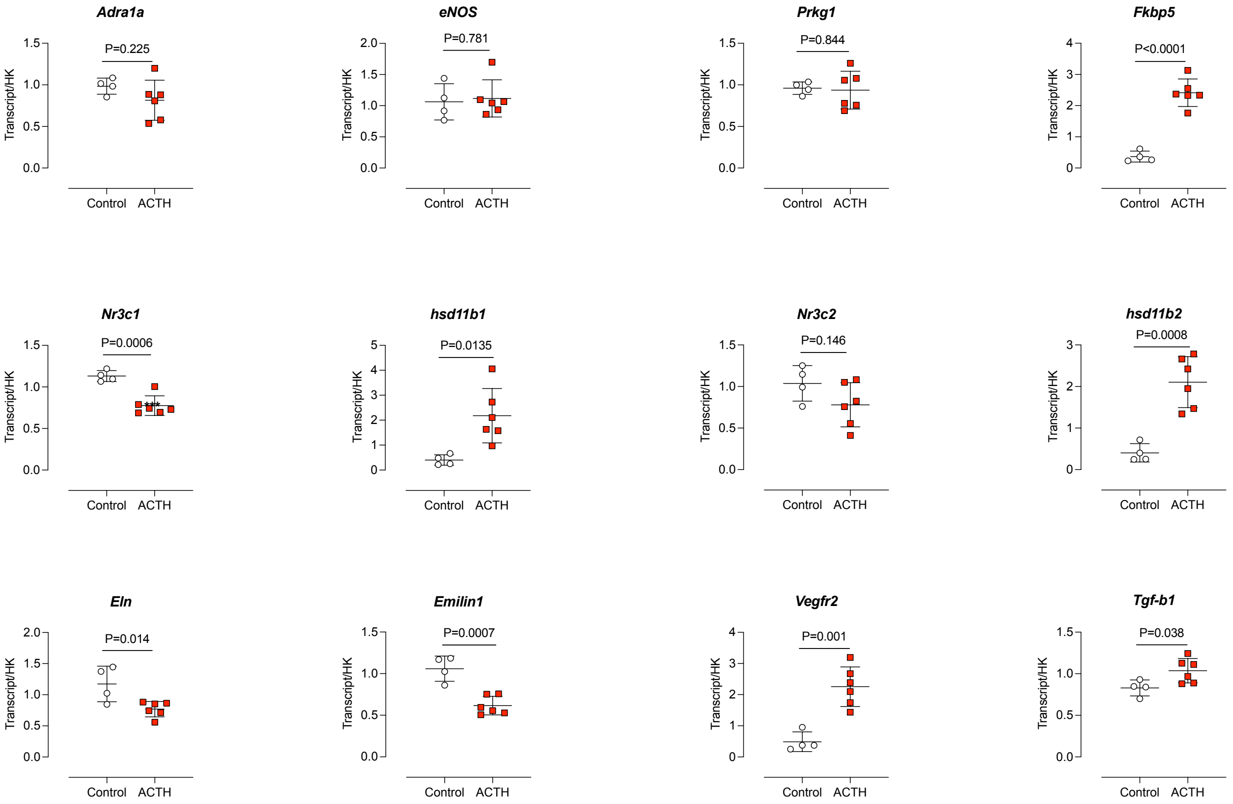

**Figure S5**. Mice were infused with ACTH (red square; n=8) or saline vehicle (controls; open circle; n=6) for 14 days and second order mesenteric artery segments were isolated. Artery rings were mounted on a wire myograph for measurement of A) the contractile response to phenylephrine, expressed as a % of the maximal constriction induced by 125 mmol/l KCL; the dilation induced by B) acetylcholine (endothelium-dependent) and C) sodium nitroprusside (endothelium-independent). Data are group mean ±SD and statistical comparisons were made by two-way ANOVA for the main effects of concentration, treatment group and interaction. No significant differences were found.

**A) Response to phenylephrine**

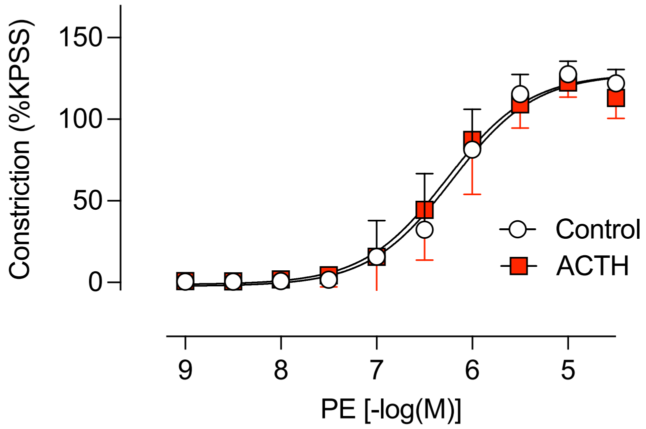

**B) Response to acetylcholine**

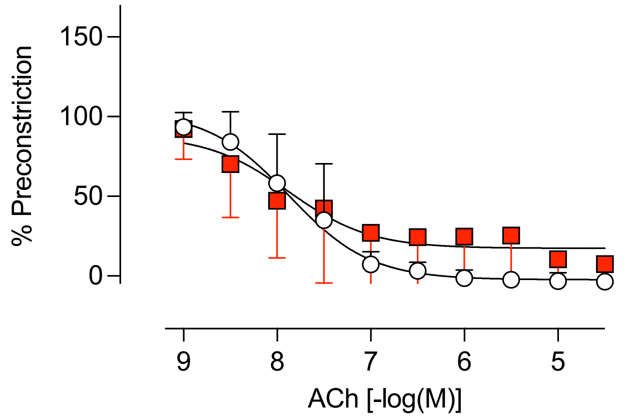

**C) Response to sodium nitroprusside**

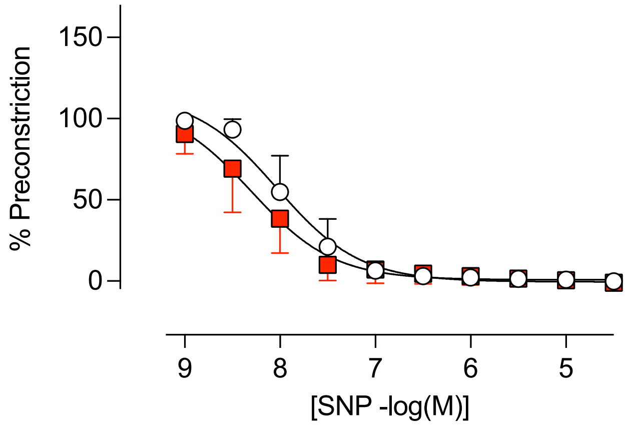

**Figure S6**. mRNA abundance of different genes in the second order mesenteric artery isolated from mice treated for 14 days with vehicle control (n=4; open symbols) or ACTH (n=6; red circles). Target genes were *Adra1a* (encoding α1A adrenoreceptor), *Nos3* (endothelial nitric oxide synthase), *Prkg1* (encoding α- and β- isoforms of soluble cyclic GMP-dependent protein kinase 1), *Fkbp5* (FK506 binding protein 5), *Nr3c1* (glucocorticoid receptor), *Hsd11b1* (11β-hydroxysteroid dehydrogenase type 1), *Nr3c2* (mineralocorticoid receptor), *Hsd11b2* (11β-hydroxysteroid dehydrogenase type 2), *Eln* (elastin), *Emilin-1*, *Vegfr2* and *Tgf-b1.* A panel of appropriate reference (housekeeper; HK) genes, *Hprt*, *Rn18s*, *Actb* and *Tbp*, which did not differ between treatment groups. Data were log transformed to ensure symmetry of the data set and normality tests were conducted. Individual data points are shown, with group mean±SD. Comparisons were made with unpaired t-test; P-values as shown.

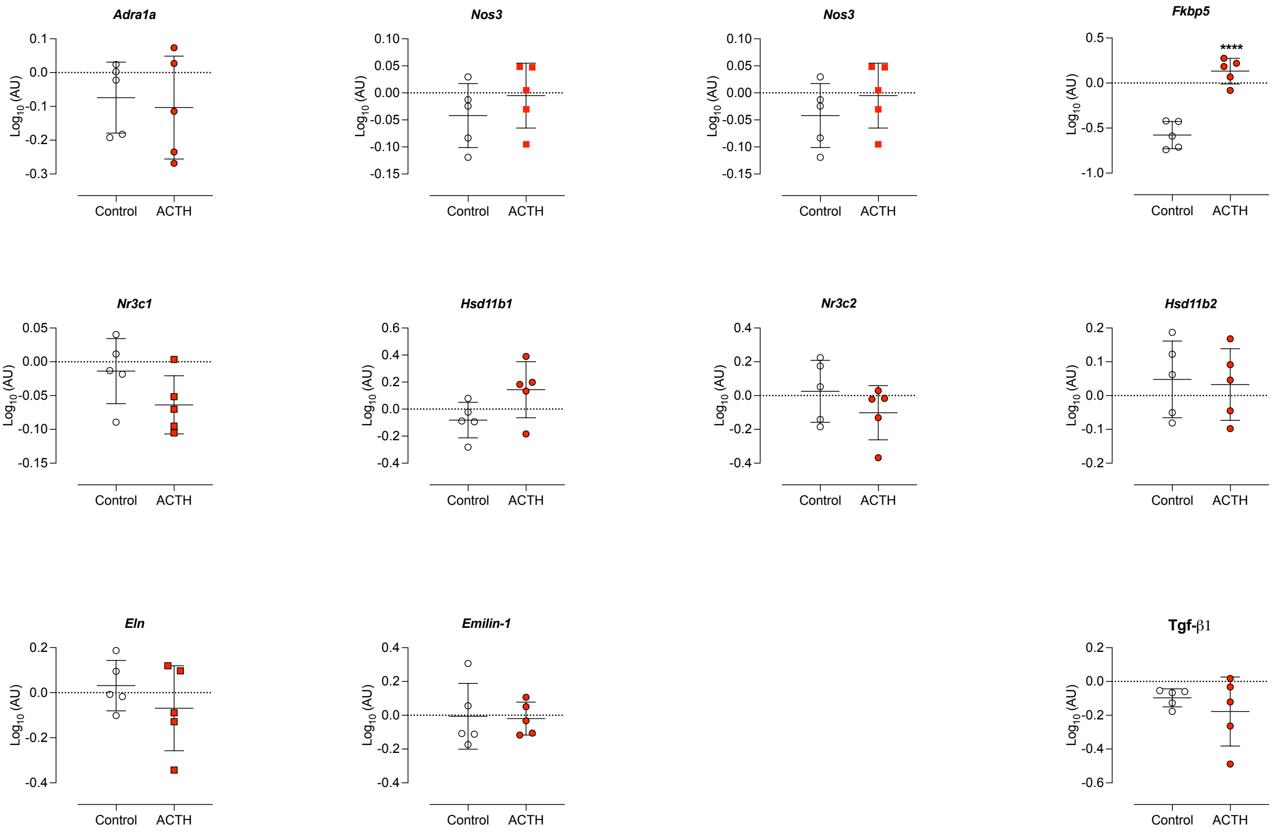

**Figure S7**. *Ad libitum* drinking behaviour from a bottle containing 1.5% NaCl was measured in n=8 mice before and after 14 days of ACTH infusion. Individual data points and group mean±SD are shown. Statistical comparisons were by paired t-test, with P-value shown.

**A) Self-directed intake of 1.5% Saline**

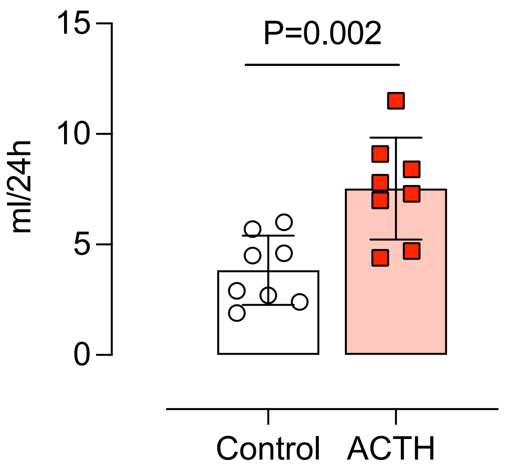

**Figure S8**. Plasma A) corticosterone and B) aldosterone in mice on a control diet (white circles), high salt diet (grey circles), ACTH + control diet (red squares) and ACTH + high salt diet (Black Squares). Corticosterone was measured in tail blood samples taken from conscious mice at 7am and 7pm local time. Aldosterone was measured in trunk blood. Individual data points are shown and comparisons were by ANOVA with Holm-Šidák for planned comparisons. P-values as shown.

**A) Plasma corticosterone**

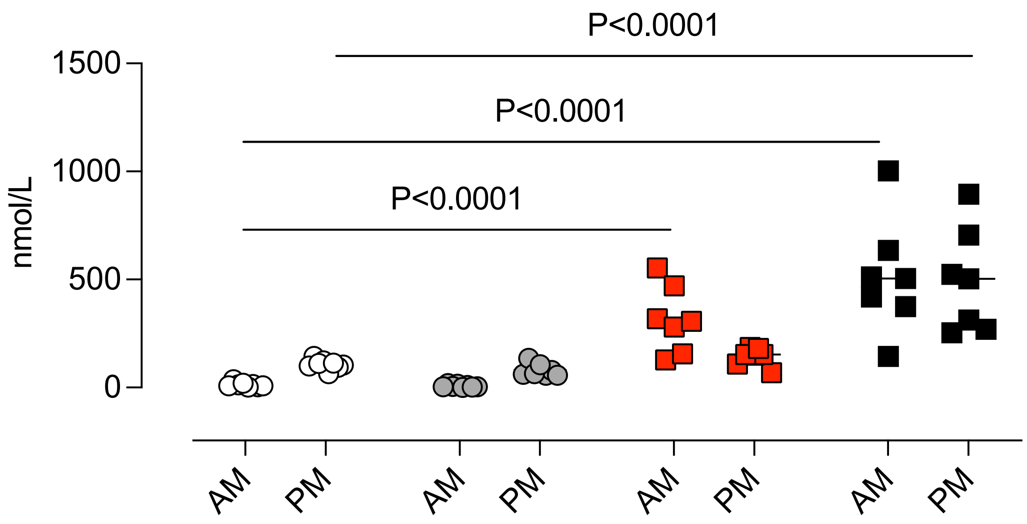

**B) Plasma aldosterone**

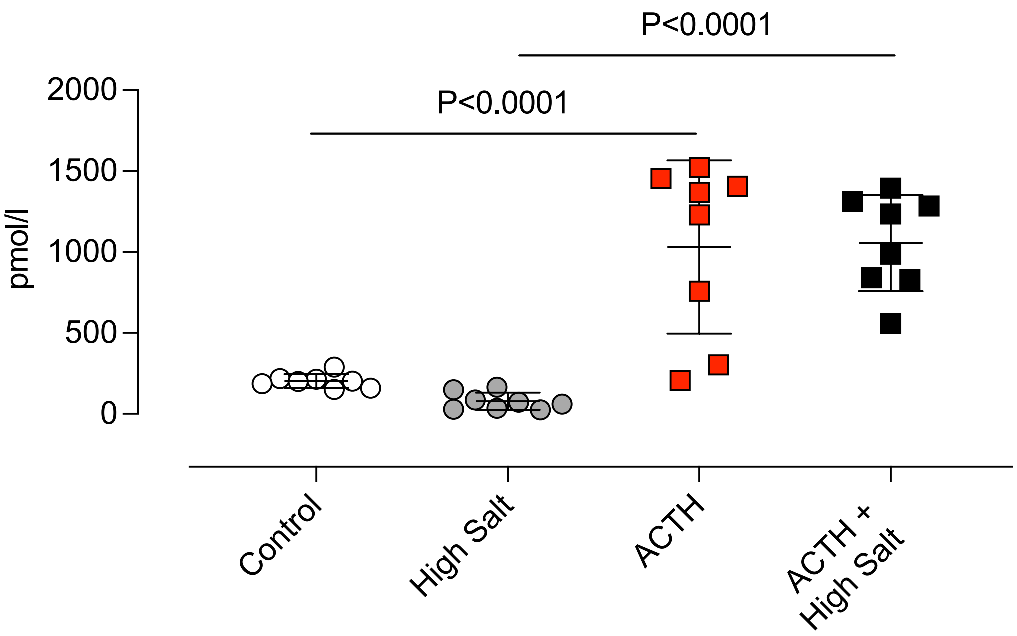
